# Oscillatory dynamics of sustained attention states

**DOI:** 10.1101/2024.09.25.614991

**Authors:** Rodolfo Solís-Vivanco, Louise Barne, Anthony Harris, Xin Liu, Nilli Lavie

## Abstract

Sustained attention allows concentration on a task over long periods of time. This ability fluctuates, with periods of effective focus (‘in-the-zone’) and periods of increased performance variability and susceptibility to errors (‘out-of-the-zone’). Little is known about the neural dynamics underlying these states and their fluctuations during sustained attention tasks. To address this, we had thirty young adults perform the gradual onset continuous performance task (gradCPT), during which their EEG and responses were recorded. States of sustained attention (out-vs. in-the-zone) were identified based on the variance time course of participants’ RT. Out-of-the-zone states were associated with increased errors of commission and reduced perceptual sensitivity compared to in-the-zone states, as expected. Importantly, a significant decline in theta oscillations at mid-prefrontal regions was found during out-of-the-zone (vs. in-the-zone) states over a ∼400 ms period around the transition point between stimuli, and the extent of this decline predicted commission errors and response bias. In addition, individual differences in the variability of midfrontal theta along the task were associated with RT variability. Finally, participants exhibiting greater theta variability showed a more pronounced decline in perceptual sensitivity when being out-of-the-zone and less stable RTs compared to those with lower variability. Our results suggest that states of diminished sustained attention, even during short lapses, are characterized by a reduction in midfrontal theta activity, and that fluctuations in this rhythm covary with fluctuations in attentional control.

## INTRODUCTION

Sustained attention, namely the ability to maintain attention focus throughout a mental task, is critical for effective information processing and performance (Esterman et al. 2019). Sustained attention is known to decline with a longer time-on-task (Gilsoul et al. 2024), resulting in increased RT and error rates, typically explained by increased “mental fatigue” and decreased vigilance (Warm, Parasuraman, & Matthews, 2008; Wascher, et al., 2014). Cognitive control also deteriorates with longer time-on-task, as evidenced in increased failures to inhibit responses to designated ‘NoGo’ target stimuli in ‘Go/NoGo’ paradigms (Manly, Robertson, Galloway, & Hawkins, 1999; Terentjeviene, et al., 2018).

In addition to the decline with time-on-task, the ability to sustain attention focus is also found to fluctuate throughout the task period. A greater level of attention fluctuations is reflected in increased response variability, along with an increase in response errors and failures to inhibit a dominant ‘Go’ response (Esterman, Noonan, Rosenberg, & Degutis, 2013; Flehmig, Steinborn, Langner, Scholz, & Westhoff, 2007; Rosenberg, Noonan, DeGutis, & Esterman, 2013). Increased response variability has been further established as a senstive marker of deficits in sustained attention abilities either due to immature ability in earlier stages of development (Hobbiss & Lavie, 2024) or to clinical deficits (Epstein, et al., 2011; Tamm, et al., 2012). A growing body of studies has focused on characterizing the fluctuations of sustained attention states throughout the task, highlighting that periods of effective attention focus (‘in-the-zone’ states) are interspersed with periods of reduced attention focus (‘out-of-the-zone’ states), that are defined by increased response variability, and are associated with reduced detection performance and increased response inhibition failures, compared to in-the-zone states (Fortenbaugh, DeGutis, & Esterman, 2017; Jun & Lee, 2021; Rosenberg, Noonan, DeGutis, & Esterman, 2013). These attention states have been identified during a modified continuous performance task (CPT), which maintains the demands on response inhibition in a minority of trials (defined as the NoGo trials) but replaces the abrupt onset of each trial with a gradual change from one image to the next, hence termed gradual-onset CPT (gradCPT, (Rosenberg et al. 2013)). Since abrupt onsets are known to capture attention and can thus facilitate task focus in a bottom-up manner (reducing the demand on cognitive control), their removal in the gradCPT makes this task particularly sensitive to detect periodic states of reduced attention focus (Esterman, Noonan, Rosenberg, & Degutis, 2013; Fortenbaugh, Rothlein, McGlinchey, DeGutis, & Esterman, 2018). Indeed, research employing this task has shown that, although the frequency of out-of-the-zone states increases with time-on-task, as would be expected from increased demands on sustained attention, the task is sensitive to reveal out-of-the-zone states in early task periods too (Esterman, Noonan, Rosenberg, & Degutis, 2013).

So far, cognitive neuroscience research of sustained attention states (out-vs. in-the-zone) has typically used fMRI and revealed the involvement of the Default Mode Network (DMN) and the dorsal attention network (DAN) in these states (Esterman et al. 2013; Fortenbaugh et al. 2018; Kucyi et al. 2016, 2017). In contrast, EEG research has not yet examined the oscillatory dynamics of the attention zone states that reflect a change in the ability to sustain a consistent level of performance (as in ‘in-the-zone’ states) or states of being ‘out-of-the-zone’ which are typified by increased RT variability and reduced detection accuracy as they fluctuate throughout a continuous performance task. It thus remains unclear whether out-versus in-the-zone states involve fine-grain changes in neurophysiological dynamics, and whether these might correspond to current views of the oscillatory correlates of sustained attention. Specifically, EEG research has suggested particular roles in sustained attention for oscillations in the frequency bands of theta (4-7 Hz) and alpha (8-14 Hz). In the case of theta oscillations, an increase in power over frontal midline channels is typically found and is thought to reflect increased demand for cognitive control to counteract the reduction in attention capacity with time-on-task, as revealed by increased lapses of attention and therefore increased number of errors and longer RT (Cavanagh and Frank 2014; Umemoto et al. 2019; Wascher et al. 2014). For alpha oscillations, increased power in frontal and posterior brain regions is thought to reflect reduced cortical excitability associated with declined vigilance with longer time-on-task (see Clayton et al. (2015) for a review and Tran et al. (2020) for a meta-analysis). Interestingly, a recent study demonstrated that cortical excitability as indicated by alpha power in frontal, temporal, and parietal cortex can also wax and wane throughout the task in a slow regular rhythmic pattern (∼.07 Hz), and this is coupled with corresponding changes of entrainment to sounds (being lower when alpha power is higher), thus suggesting slow rhythmic fluctuations may be involved in the application of sustained attention to external stimuli (Kasten et al. 2024).

Regular fluctuations of cortical excitability as measured by neural oscillations in a high frequency band (HFB) in frontal and parietal channels at cued location-related electrodes have also been found to nest within a theta rhythm pattern power during cue-to-target periods (500 – 1700 ms) of spatial cueing tasks (Helfrich et al. (2018). A correlation between the phase of theta (reflecting an either high or low level of cortical excitability) and target detection supports that rhythmic attention sampling is involved in spatial attention tasks (Harris et al. 2018; Landau et al. 2015; Landau and Fries 2012). However, it is not clear whether the rhythmic sampling at the theta band in these tasks reflects a general mechanism of sustained attention or instead reflects the specific nature of location sampling in tasks involving spatial orienting of attention. One could argue that sustaining attention focus on a specific location in the field is particularly demanding on the fronto-parietal network underlying attention orienting in space. Indeed, Raposo et al. (2023) demonstrated a causal role for oscillatory activity (theta and alpha/low beta) in the fronto-parietal network in attentional sampling in patients with unilateral fronto-parietal lesions who performed a lateralized spatial attention task similar to that used in Helfrich et al. (2018).

A general mechanism of sustained attention that imposes no demand on spatial orienting (e.g. presenting stimuli in the same fixed location throughout the task as is typical in most CPT and sustained attention to response tasks (SART)), may recruit different networks with potentially different oscillatory dynamics. For example midfrontal theta, reflecting the involvement of a generalized cognitive control attention mechanism, as well as alpha band oscillations that may reflect, at least in part, the level of cortical excitability in the regions involved in the processing of task stimuli (e.g., occipital cortex for visual tasks (Wolfgang Klimesch 2012; Stephani et al. 2021) and temporal auditory cortex regions in relation to auditory stimuli (Kasten et al. 2024)), along with changes of activity in frontal top-down control mechanisms.

In addition, the variability of theta power may also play a role, since much like RT variability is thought to reflect an unstable ability to sustain attention focus (Castellanos and Tannock 2002; Machida et al. 2022), the variability of an oscillatory signal that is thought to mediate attention may also reflect irregular fluctuations in this process (e.g. midfrontal theta power variability may reflect fluctuations of attentional monitoring during the task performance, which lead to states of increased RT variability).

Consistent with this suggestion, Wang et al. (2020) examined whether variability of frontal theta power correlated with RT variability (measured with SD) throughout task performance in a forced choice RT task (requiring pressing one of two keys to indicate the direction of an arrow). No correlation was found between the variability of theta power and RT variability. However, the task used in their study involved abrupt display onsets and responses being made on each trial, and these factors can contribute to maintaining attention on the task. It is thus plausible that their method was not sensitive to reveal fluctuations of sustained attention states. We note that the RT variability in terms of fluctuations of amplitude of low-frequencies (0.01–0.027 Hz) showed a significant correlation with theta activity in one left frontal channel. However, as these findings did not relate to decrement in task performance, their functional significance is unclear.

A clearer finding that did relate the variability of the theta phase alignment to behavioural performance was found in Reteig et al. (2019). They found that reduced levels of sustained attention with longer time-on-task, as measured with target detection sensitivity, could be predicted from the consistency of phase alignment across trials (i.e. the inter-trial phase clustering, ITPC) in the theta band signal, such that longer time-on-task led to both lower ITPC theta and lower detection sensitivity. Moreover, these changes were correlated, such that the reduced detection sensitivity with longer time-on-task could be predicted from the effect of time-on-task on theta ITPC. These findings strongly suggest that the effects of longer time-on-task (within an 80 min task) on sustained attention are mediated by reduced ITPC of theta oscillations. However, it remains unclear whether theta phase alignment can reflect fluctuations of short-term states of sustained attention (e.g. in-vs. out-of-the-zone) in task paradigms that are more sensitive to reveal variability in these states throughout the task period (including early task periods).

Overall then, the oscillatory dynamics associated with sustained attention states of being in-the-zone to being out-of-the-zone, as revealed by increased response variability throughout a sustained attention task, have not as yet been identified. In addition, any potential role for the oscillatory correlates of these states and the variability of power in predicting individual differences in sustained attention, remains to be established as well. We thus set out to investigate these questions in the present study by using a sustained attention paradigm that is well established to discern sustained attention states (the gradCPT) and examined the oscillatory correlates of in-the-zone and out-of-the-zone states in this task. Considering that the brain regions that give rise to relevant oscillations for attention performance seem to vary depending on specific task demands (e.g. for theta: midfrontal regions for cognitive control (Clayton et al. 2015) or fronto-parietal regions for spatial attention (Helfrich et al. 2018; Raposo et al. 2023)), we added a source analysis to identify the likely cortical sources of our EEG findings. We also investigated the power variability of any oscillations that were found to be associated with sustained attention states and assessed whether either oscillatory power or power variability underlying different attentional zone states can predict behavioural performance indicators of sustained attention.

Based on previous research of the oscillatory dynamics of sustained attention and the evidence showing increased RT variability as a marker of reduced sustained attention (Chidharom et al. 2021; Flehmig et al. 2007), our main hypotheses were that both the power and power variability of theta or alpha activity would reflect the distinct states of attentional zone (out-versus in-the-zone). Moreover, we expected that individual differences in both of these potential neural markers of sustained attention would predict attentional performance as revealed by behavioural outcomes. We also added an exploratory analysis of power and power variability in the beta frequency band (15-30 Hz), since to the best of our knowledge our study is the first to uncover the oscillatory dynamics of sustained attention in the widely used gradCPT paradigm and these may draw also on neural mechanisms associated with beta power (perhaps in a similar manner to an increase in beta power associated with improved sustained attention in previous studies using the traditional CPT paradigm (Riccio 2002). We therefore examined whether beta power and its variability would also be involved in predicting sustained attention states in our task. Finally, we also considered ITPC in relation to sustained attention states, since the phase alignment of neuronal oscillations has been shown to play an important role in the fine temporal attunement of attentional and cognitive control and has been associated with levels of sustained attention (Reteig et al. 2019).

## METHODS

### Participants

Thirty-one adults were recruited via the University College London SONA system. This sample size allowed a power of 0.85 to detect effect sizes of 0.5 and above in paired samples t-tests (comparisons of in-vs. out-of-the-zone states). One participant was excluded from the analysis due to difficulties recording their EEG data. The final sample was of 30 participants (mean age=25±5.1, 15 women, 26 right-handed). All participants reported normal vision and no clinical (neurological or mental health) diagnosis. The study was approved by the UCL Research Ethics Committee, and all participants gave written informed consent before any experimental procedure took place. Participants were compensated £20 for taking part in the study.

### Experimental task

The gradCPT task was administered. Similar to (Esterman et al. 2013), the task stimuli included 10 round, grayscale photographs of mountain scenes and 10 city scenes of 8.9 cm circumference, which at a fixed viewing distance of 64 cm (held with a head rest) subtended 8 degrees of visual angle. All the images were displayed against a gray background on a gamma-corrected Dell monitor with 2560 × 1440 resolution and refresh rate set to 60 Hz. The images were calibrated to have a mean luminance matching the gray background and 20% Michelson contrast. The images consisted of city scenes (90% of the trials) or mountain scenes (10% of the trials), with a gradual transition from one to the other during an 800 ms period (which we henceforth term trial, though with the gradual transition between scenes, the presentation appearance was of a continuous stream, rather than discrete trial-by-trial onsets). Participants were instructed to respond by pressing the space bar on a standard computer keyboard, using their dominant hand, each time they detected a city scene (Go trials) and withhold their response for mountain scenes (NoGo trials). Each frame displayed a composite of two scenes, with different levels of coherence for each of the scenes starting with 0% coherence of the trial scene (scene Nth) and 100% coherence of the previous trial scene (scene Nth-1) and building up gradually up to 100% coherence of the trial scene and 0% of the subsequent trial scene (Nth+1). Transition point is referred as the starting point of a subsequent scene presentation (0% coherence). To achieve a gradual transition between the scene composites, a linear pixel-by-pixel interpolation (in steps of 2.08% coherence change per frame) was used (Figure 1). Trials were presented in random order with the constraint that no scene was repeated on consecutive trials. Participants were instructed to keep their gaze on a white fixation point in the middle of the images throughout the experiment.

**Figure 1.**
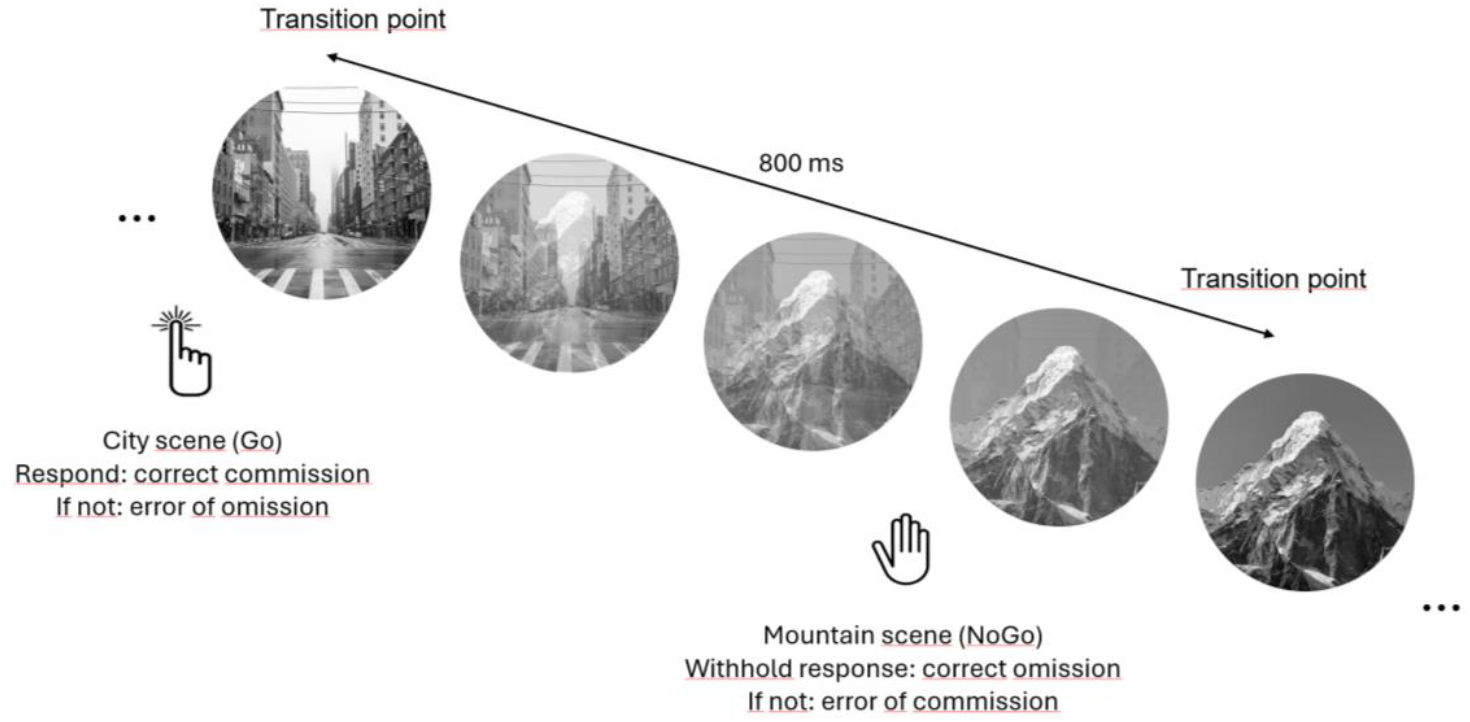
GradCPT task procedure. Grayscale images of city (Go) and mountain (NoGo) scenes were presented, with a smooth transition from one image to the next one, through varied level of coherence for each scene in composite images of the trial scene mixed with the previous trial scene. The figure depicts only 5 out of several frames within each 800 ms trial period and exemplifies a city trial shown on the left of the figure, starting from a transition point in which the previous trial city scene (Nth-1) is shown at 100% coherence and the current one (Nth, mountain scene) at 0% coherence, building up to 100% at end of the next trial (next transition point).

### Procedure

After signing the informed consent, participants underwent 2 practice blocks. The first one was presented at ∼ 50% of the normal speed for 30 seconds. The second one was presented at normal speed for 60 seconds. After practice, each participant completed a total of 2250 trials made up of 5 blocks with 6 minutes duration each. Participants were forced to take at least a 30 seconds break between each block. The total duration of this experiment was approximately 40 minutes. Participants were encouraged to respond as quickly and accurately as possible.

### EEG recording and pre-processing

We recorded a digital EEG while participants performed the gradCPT in a dimly lit room. We used a BioSemi Active Two system with a sampling rate of 1024 Hz and offsets between +/-20 along the recording. The EEG recording was obtained from 57 silver-chloride electrodes placed according to the international 10/10 system. In parallel, we recorded the pupillary dilation of both eyes during task execution with an Eyelink 1000 eye tracker (reported in (Liu 2019)). Seven electrodes (F7, AF7, FP1, FPZ, FP2, AF8, F8) were excluded from the standard 64 channels due to the use of a forehead rest to stabilize the participants’ heads, as required by the eye tracking system. The Common Mode Sense (CMS) and Driven Right Leg (DRL) electrodes were employed as grounds. All scalp electrodes were referenced to the CMS during recording.

All offline EEG analyses were conducted using custom MATLAB 2017b scripts and the Fieldtrip toolbox (Oostenveld et al. 2011). The EEG was down sampled to 500 Hz and segmented in 1600 ms windows, with -800 ms and +800 ms relative to the transition point between stimuli (100% coherence of the Nth-1 trial scene and 0% coherence of the Nth trial scene, Figure 1). The EEG segments were re-referenced to the average of all EEG channels and baseline-corrected for the entire segment. Subsequently, we removed noisy or atypical segments in terms of voltage variance using the ft_rejectvisual function. The remaining segments were visually inspected using the ft_databrowser function to eliminate those with noise or artifacts, including blinks and muscle-related artifacts. For all segments, we identified RT, response type, and falling out-or in-the-zone (as described below). The mean number of selected trials for being out- and in-the-zone was 501±95 and 1249±175, respectively. Before the task, we recorded 3 minutes of resting-state EEG while participants kept their eyes closed, whose results will be reported elsewhere.

### Behavioural analysis

We recorded responses and calculated accuracy and error rates, as well as RTs from the transition point of Go trials (100% coherence of the previous trial scene and 0% of the current one, see Figure 1). Following the protocol in (Esterman et al. 2013), trials with deviant RTs (before 70% coherence of the previous image and after 40% coherence of the current scene) or with more than one button presses (ambiguous presses) were treated as follows. Presses were assigned to an adjacent trial if 1 of the 2 had no response. If both adjacent trials had no response, the press was assigned to the closest trial, unless one was a NoGo scene, in which case it was assumed that participants correctly responded to the Go scene. Finally, if there were multiple presses that could be assigned to any 1 trial, the fastest response was selected.

Performance was evaluated in terms of mean RT, variability of responses measured by the coefficient of variation (CoV = SD/M), commission response rates for both Go (i.e. correct commissions) and NoGo (i.e. errors of commission) trials, as well as the non-parametric estimate (A’) of discrimination sensitivity index (Zhang & Mueller, 2005) and response criterion (β).

Attention zones (out- and in-) were defined using a variance time course (VTC) analysis (Esterman et al. 2013), which consists of calculating the absolute z-scores of RTs in Go trials. The sliding window approach of averaging VTC values over nine consecutive trials (7.2 s) was applied to extract accuracy measures during the attention zones. Trials in-the-zone were those whose RTs were within 1 SD with respect to the mean, while out-of-the-zone were those either over 1 SD or below -1 SD.

### EEG analysis

The EEG data were explored through time-frequency (TF) representations of power and phase alignment across trials. All TF analyses were centered around the transition point between stimuli (100% coherence of the Nth-1 trial scene and 0% coherence of the Nth trial scene) and included 3 to 40 Hz oscillations, therefore encompassing theta (3-7 Hz), alpha (8-14 Hz), and beta (15-40 Hz) bands. Only segments centered around correct commissions (Go trials) were further analyzed. Power was calculated through a fast Fourier transformation (FFT) along an adaptive sliding time window including 2 cycles of each oscillation (ΔT = 2/f; e.g. 200 ms for 10 Hz). A Hanning taper (also ΔT long) was multiplied by the data before the FFT. All power values were log-transformed and averaged across trials regarding their evaluated condition (i.e., attention zones). The phase stability across trials was explored using the phase data from the FFT and calculating the the Inter-trial Phase Clustering (ITPC), also known as Phase Locking Factor (PLF) (Tallon-Baudry et al. 1996). The ITPC over N trials is defined as:

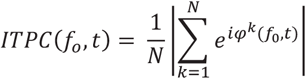

where *φ*^*k*^*(f*_*0*_, *t)* corresponds to the estimated phase at frequency *f*_*0*_ and time t resulting from the time-frequency analysis. The ITPC quantifies the degree to which the distribution of phase angles across trials at each time-frequency-sensor point is not uniformly distributed. An ITPC value approaching 0 suggests high phase variability, while an ITPC value of 1 indicates that all trials exhibit the same phase at a particular frequency and time point. Similar to the TF representation analysis of power, the ITPC was computed using a sliding time window that was 2 cycles in length, and a Hanning taper was applied to it. For the ITPC calculation, we randomly selected the same number of trials for being in-the-zone with respect to being out-of-the-zone. This was done because the ITPC is sensitive to the number of observations (Cohen 2014) and comparisons between zones were planned.

### Source analysis

The significant power effects (see results) were described from a source perspective as well. We used the dynamic imaging of coherent sources (DICS) technique, a frequency-domain beamforming approach, to estimate the absolute power at the source level in the entire brain. The cross-spectral density matrices were obtained by applying a multitaper frequency transformation approach on data measured from the channels. For each participant, a realistic description of the brain was constructed based on an adult template MRI provided by the Fieldtrip toolbox and segmented in grey matter, white matter, and cerebrospinal fluid (Oostenveld et al. 2011). After masking for the grey matter tissue only, a head model was created using a forward solution based on three concentric spheres, which was then used to create a source head model based on a grid with a 1 cm resolution. The lead field and the cross-spectral density were used to calculate a spatial filter for each grid point and the spatial distribution of power was estimated for each zone state (in- and out-) in each participant. A common filter was used when comparing the zone states (based on the cross-spectral density matrices of the combined states). The estimated power was averaged over trials and log-transformed. The power difference between zone states was calculated and averaged across participants. The source estimates were plotted on a standard MRI brain found in SPM8.

### Statistical analyses

For behavioural data, paired t-tests were performed to evaluate the effect of time-on-task within a continuous task-block (0-3 minutes vs. 3-6 minutes) on performance measures and on the out-of-the-zone occurrence rate. We also ran comparisons between attention zones (out-vs. in-the-zone) on accuracy and detection sensitivity (A’), and response criterion (β) using paired t-tests. For EEG data, we first compared the time-frequency representations of power and ITPC between out-versus in-the-zone periods using a cluster-based nonparametric randomization test (Maris and Oostenveld 2007). This test manages the control of the Type I error rate in scenarios where there are multiple comparisons involving sensors, frequencies, and time periods. It does so by grouping together neighboring sensors, time points, and frequency points that exhibit a similar effect. We examined frequencies ranging from 3 to 40 Hz, incrementing by 1 Hz, therefore including theta, alpha, and beta oscillations. This analysis was performed from -800 ms to +800 ms with respect to the transition point between stimuli (100% coherence of the Nth-1 trial scene and 0% coherence of the Nth trial scene). Electrodes that exhibited a t value difference between conditions exceeding a pre-established threshold (p <.05) were chosen and subsequently clustered based on their spatial proximity. The sum of the t values within a cluster served as the cluster-level statistic. The cluster with the highest cumulative t values was used as the test statistic. To determine the statistical significance of a particular effect, we generated a reference distribution by randomly permuting the data between the two conditions and recalculating the test statistic 1000 times (Monte Carlo estimation). Effect sizes for these tests were calculated from the average over the cluster as detailed in the Fieldtrip website (https://www.fieldtriptoolbox.org/example/effectsize/). To determine whether EEG changes between out-versus in-the-zone were associated with task performance differences among participants, we calculated the magnitude of power change between zones (out-minus in-) in time-frequency-channels of interest (as revealed by the nonparametric test, see Power section of Results) and ran correlation analyses (Pearson (r) or Spearman (rho) depending on the distribution of the data for each variables) with RT CoV, commission responses rate, sensitivity (A’), and response criterion (β).

We also assessed whether theta or beta power and power variability and their differences between zones (out-minus in-) were predictive of differences in task performance among participants, as follows. Based on our non-parametric analysis that identified time-frequency-channel clusters of significant differences between out-vs. in-the-zone periods in different frequency bands (i.e. one identified cluster for theta and two clusters for beta along the -/+ 800 ms time windows (see Power section of Results and Figures 3B and 3D)), for each trial we averaged the power values within the time-frequency-channels comprising each cluster. Specifically, for theta (4-7 Hz) power it was averaged within 400 ms around the transition point at frontal-midline channels, and for beta, it was averaged independently within two time windows: 200 ms around the transition point and from 300-500 ms afterwards, both at left central channels. The associations were then assessed between task performance for power difference between zones (out-in) for each of these three clusters and the behavioural measures of RT CoV, correct commissions rate, errors of commission rate, sensitivity (A’), and response criterion through Pearson (r) or Spearman (rho) correlation analyses. The same analyses were also run for overall power CoV (regardless of state zone) in both frequency bands and behavioural measures.

Finally, to further explore whether individual differences in theta and beta power changes between zones and theta and beta power CoV were associated with differences in task performance, participants were divided by a median split based either on power change (low/high) between zones or overall power CoV (low/high), independently for theta and beta. At first, we compared in those groups task performance changes in sensitivity (A’) and response criterion (β) between the attention zone states (out-vs. in-the-zone) with independent samples t tests. Additional analyses were also run with mixed model ANOVAs with the between-subject factor of group (i.e. low vs. high power change or high vs. low overall power CoV for each frequency band) and the within-subject factors of time-on-task (0-3/3-6 min) and block number (1-5), on the variables of mean RT, RT CoV, correct commission rate, A’ and β. All ANOVA comparisons were adjusted by sphericity according to the Greenhouse-Geisser correction, and all post hoc analyses were done with a Bonferroni correction.

In order to examine which EEG feature was able to better predict performance across participants, we ran three independent multiple regression models taking those EEG measures that were sensitive to attentional states as regressors (i.e. theta power decreases in out-vs. in-the-zone, theta power CoV, and beta power decreases in out-vs. in-the-zone, as reported in the Results section) and the behavioural measures of correct commissions rate, correct omissions rate, and RT CoV as the dependent variable in each of the models.

Statistical significance for all analyses was established at p<.05. JASP (https://jasp-stats.org) and Matlab (MathWorks Inc.) softwares were used for all statistical analyses.

## RESULTS

### Behaviour

Across the 6-min runs, time-on-task led to slower (t(29)=-4.4, p<.001, Cohen’s d=-0.803, Figure 2A) and more variable (t(29)=-3.86, p=.001, Cohen’s d=-0.706, Figure 2B) RT, with both p values passing the Bonferroni correction for two multiple comparisons (.05/2= 0.025). Discrimination sensitivity (A’) decreased over time (t(29)=2.66, p=.01, Cohen’s d = 0.486, Figure 2C) as did the rate of correct commissions in Go trials (t(29)=4.09, p<.001, Cohen’s d=0.746, Figure 2D). These effects demonstrate the expected effect of time-on-task on sustained attention. The rate of errors of commission in NoGo trials however was unaffected (t(29)=0.02, p=.984, Cohen’s d=0.004, Figure 2E). The significant effects on errors passed the Bonferroni correction for 3 multiple comparisons (.05/3= 0.016).

**Figure 2.**
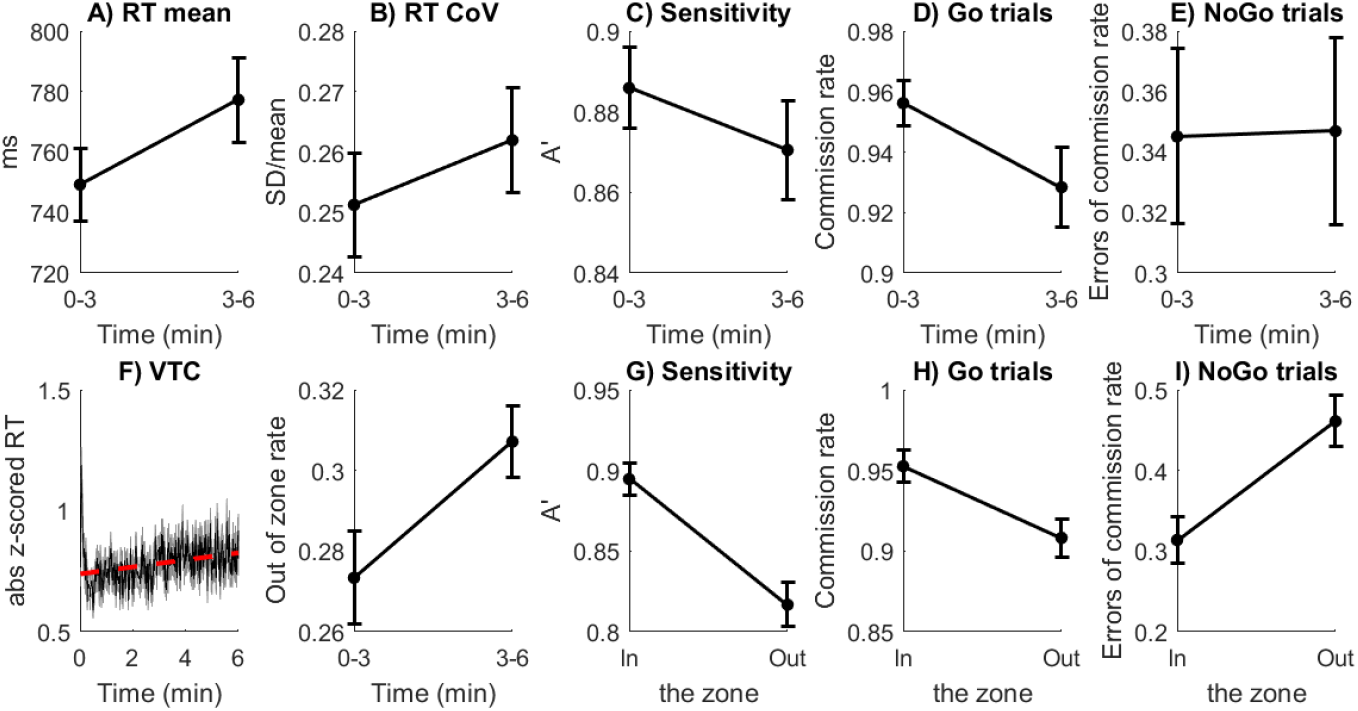
Participants’ performance (mean±SEM). With time-on-task, participants showed (A) increased mean RT to Go trials, (B) higher coefficient of variation (CoV) of Go RTs, (C) decreased discrimination sensitivity (A’), (D) lower rate of correct commissions (Go trials), and (E) no changes in the rate of errors of commission (NoGo trials). (F) Variation time course (VTC) analysis. Left panel: absolute z-scored RTs as a function of the time, and its linear trend represented by the dashed red line. Right panel: proportion of the out-of-the-zone state (VTC > 1) as a function of time-on-task. Being out-of-the-zone was associated with (G) lower discrimination sensitivity (A’), (H) lower rate of correct commissions (Go trials), and (I) higher rate of errors of commission (NoGo trials).

Considering the sustained attention states as indexed with our VTC analyses, we found that out-of-the-zone became more prevalent as a function of time-on-task (t(29)=-4.58, p<.001, Cohen’s d=- 0.836, Figure 2F, which passes the Bonferroni correction for three multiple comparisons (.05/3=0.016) when considered together with the effects of time-on-task on RT and RT variability. Out-of-the-zone states involved reduced discrimination sensitivity (t(29)=14.06, p<.001, Cohen’s d=2.567, Figure 2G), reduced rate of correct commissions (Go trials) (t(29)=14.39, p<.001, Cohen’s d=2.627, Figure 2H), and increased rate of errors of commission (NoGo trials) (t(29)=7.99, p<.001, Cohen’s d=1.458, Figure 2I) compared to in-the-zone states. These comparisons passed the Bonferroni correction for six multiple comparisons (.05/6=0.008 when considered together with the time-on-task effects on the measures of errors). Response criterion (β) was not significantly different between zone states (t(29)=1.62, p=.16, Cohen’s d =.29).

### EEG

#### Power

Figure 3A shows the results of the TF analysis of power, which revealed a decrease within the theta range (4-7 Hz) for out-versus in-the-zone states. The non-parametrical permutation test showed a significant decrease (cluster statistic=-29, p=0.002, Cohen’s d=.77) over a 400 ms period around the transition point (0 ms) of Go trials (Figure 3A, dashed rectangle). The significant effect covered a frontal-midline region, with the highest differences in F1, FC1, Fz, FC2, and FCz (Figure 3B). The beamformer source analysis revealed changes at frontal and cingular regions, with the maximum difference at medial frontal cortex (Figure 3C). Frontal theta power did not change significantly with time-on-task: it did not increase along the experiment blocks (block effect: F(1,28)=1.97, p=.11, n^2^_p_ =.07) nor changed between the first and second halves of the blocks (time effect: F(1,28)=1.66, p=.21, n^2^_p_ =.06). Thus, the decrease in frontal theta power in out-versus in-the-zone trials was specific to the states of sustained attention zones as they occurred throughout the task.

**Figure 3.**
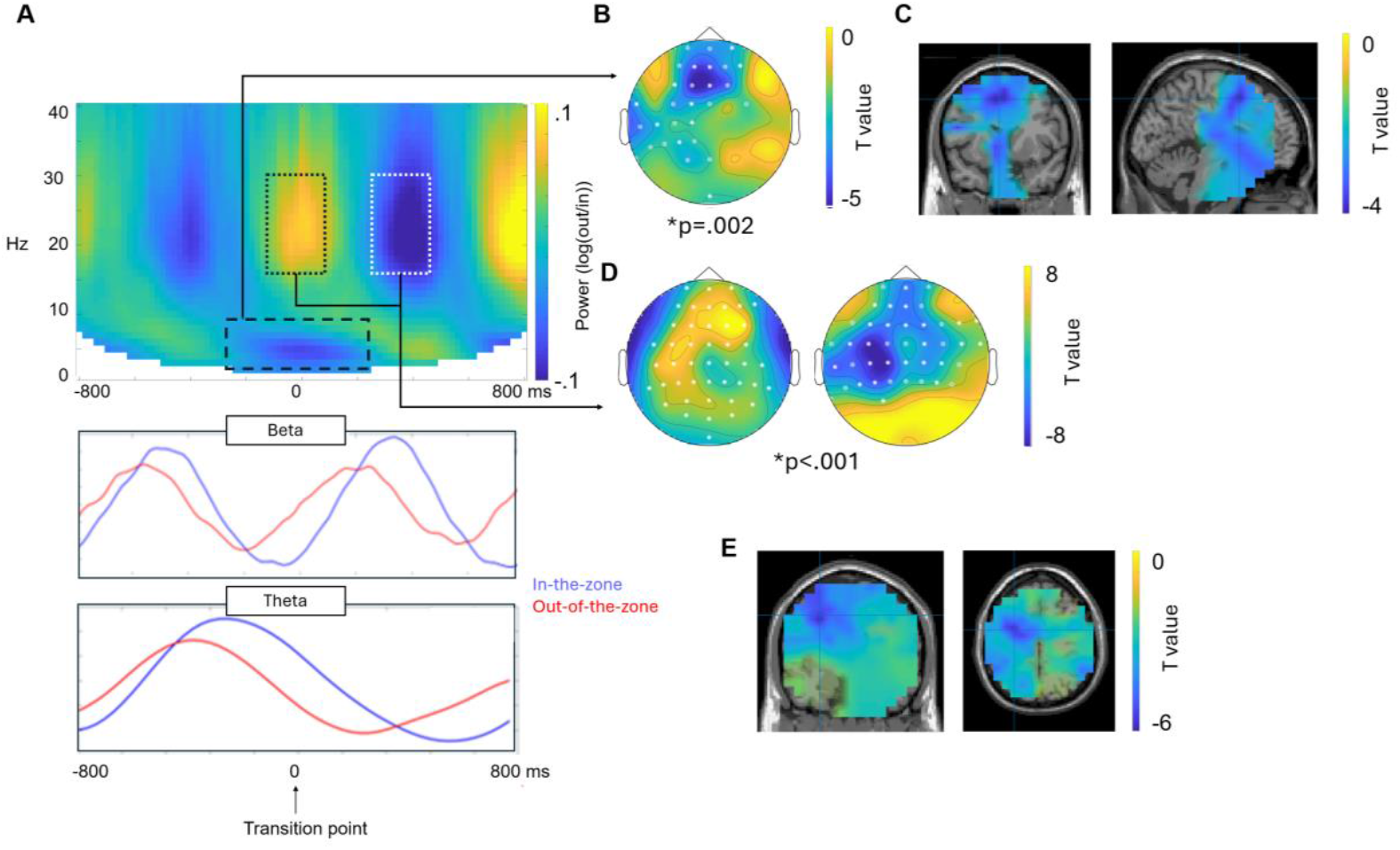
EEG power differences between out-versus in-the-zone during sustained attention. (A) Time-frequency representations of power revealed significant decreases of theta (dashed rectangle) around the stimuli transition point (100% coherence of the Nth-1 trial scene and 0% coherence of the Nth trial scene) and a fluctuating change in beta power (dotted rectangles) around the stimuli transition point and 400 ms afterwards of Go trials. The lower panels show the temporal dynamics of each band along the trial. The figures correspond to the means across channels with significant differences. (B) Statistical topography with channels where differences were significant for theta power, *p=.002. (C) Source projection on grey matter of significant differences for theta power. (D) Statistical topographies with channels where differences were significant for beta power, *p<.001. (E) Source projections of significant differences for beta power.

Figure 3A (dotted rectangles) shows that the effects of sustained attention states were also found in the beta range (15-30 Hz). The non-parametric permutation test revealed that out-versus in-the-zone states were associated with a significant increase in beta power at frontal regions around the stimulus transition point (0 ms), during Go trials (cluster statistic=45, p<.001; Cohen’s d=.97; Figure 3D) together with a significant reduction about 400 ms after stimulus transition point (cluster statistic=-42, p<.001; Cohen’s d=.94; Figure 3D), both at left central regions, around the primary motor cortex (Figure 3E).

No significant differences between zones were found in the alpha band (cluster statistics>=169, p>=0.064). Since no significant effects were found for alpha when comparing zones, no further analyses were carried out with this frequency band.

#### EEG power and differences in task performance

We further assessed whether power changes when out-versus in-the-zone were associated with differences in task performance among participants. Correlation analyses revealed that individuals manifesting a stronger decline in theta power in out-of-the-zone (vs. in-the-zone) states had higher RT CoV (r=.49, p=.006, Fisher’s z=-.53; Figure 4A) as well as higher rate of errors of commission or ‘attention lapses’ on NoGo trials (r=.45, p=.01, Fisher’s z=.48; Figure 4B) and a greater difference in attention lapses between out-compared to in-the-zone (r=.40, p=.028, Fisher’s z=.43; Figure 4C). There was also a non-significant trend for an association with a greater rate of commission responses on Go trials (rho=.33, p=.08, Fisher’s z=.34; Figure 4D), and a significant correlation with an overall increase in response bias (rho=.59, p<.001, Fisher’s z=.68; Figure 4E). There was no significant correlation between theta power decline in out-of-the zone (vs. in the-zone) states and the effect of attention zone state on the number of correct commissions in Go trials (r=.03, p=.88, Fisher’s z=.03). No correlations were found for theta power decline and sensitivity (A’, r=.14, p=.47, Fisher’s z=.14) nor with sensitivity difference between zones (rho=-.14, p=.46, Fisher’s z=-.14).

**Figure 4.**
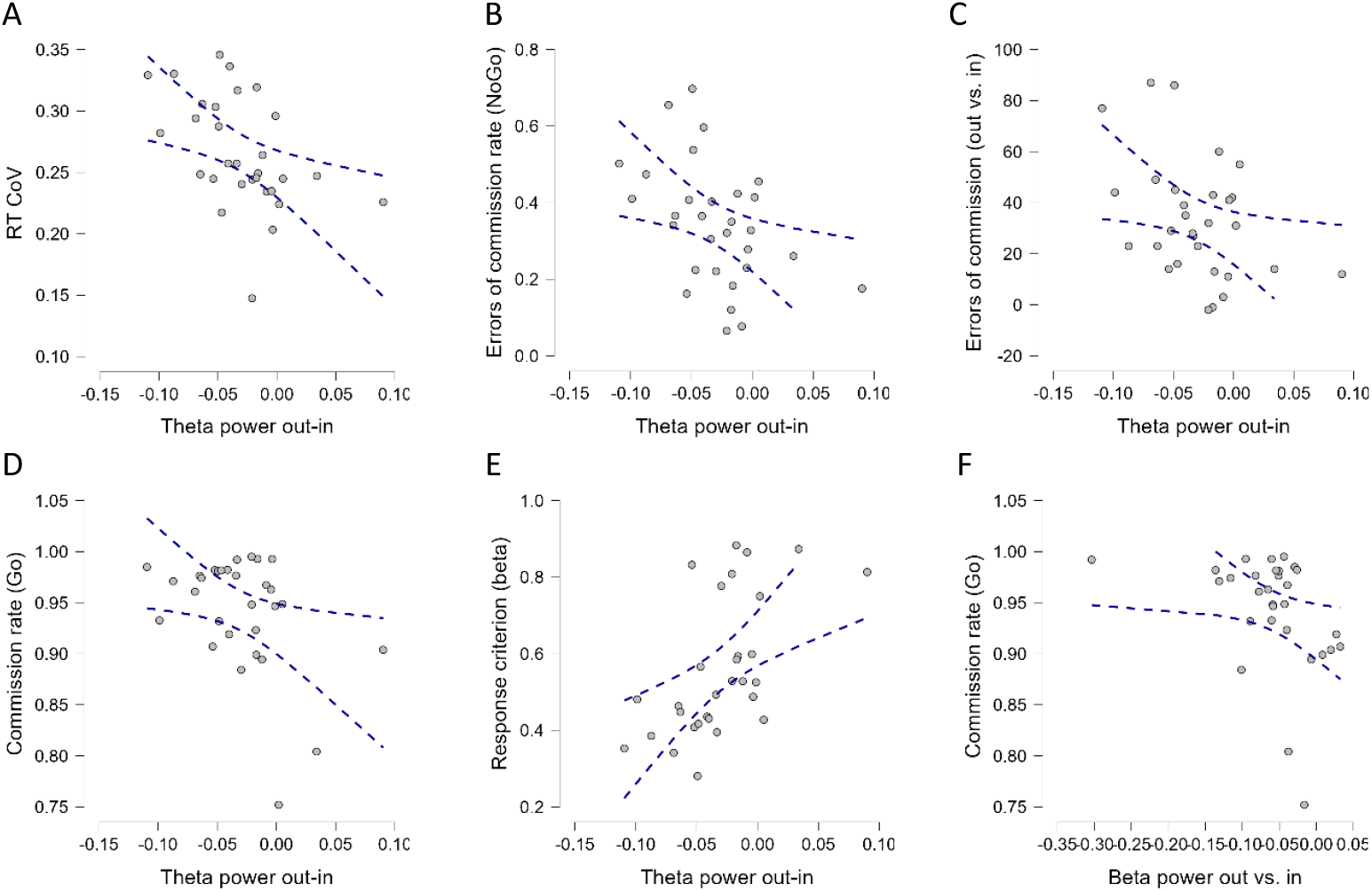
Differences in theta (A-E) and beta (F) power between attention zones (out versus in) and its association with task performance. Greater theta decline in out-of (vs. in-) the-zone is associated with (A) increased RT CoV, (B) increased (correct) commission rates in Go trials and (C) in NoGo trials (errors of commission), (D) lower response criterion, as well as (E) higher number of errors of commission in NoGo trials in out-versus in-the-zone states (F) Greater beta decline in out-of (vs. in-) the-zone was associated with an increase in the commission responses in Go trials. Blue dotted lines indicate confidence intervals (95%).

Beta power correlated with task performance, such that a stronger power decline in out-of-the-zone (compared to in-the-zone) states around 400 ms after stimuli transition point was associated with increased rate of correct commissions (Go trials) (rho=.44, p=.02, Fisher’s z=.47, Figure 4F) and increased overall response bias (rho=.39, p=.03, Fisher’s z=.41). No correlations were found between the beta power differences between zones around the transition point and behaviour (all r<.35, all p >.06).

#### EEG power variability

There was no difference between either theta or beta power CoV between out-of-the-zone and in-the-zone states, t(29)=.46, p=.65, and t(29)=.06, p=.96, respectively. However, theta power CoV was positively correlated with RT CoV (r=.38, p=.04, Fisher’s z=.39; Figure 5A). No significant correlations were found between theta power CoV and other behavioural measures (all p >.10). No significant correlations were found between beta power CoV in any of the time windows (during the transition point, or 400 ms after it) and any of the behavioural measures either (all p >.20).

**Figure 5.**
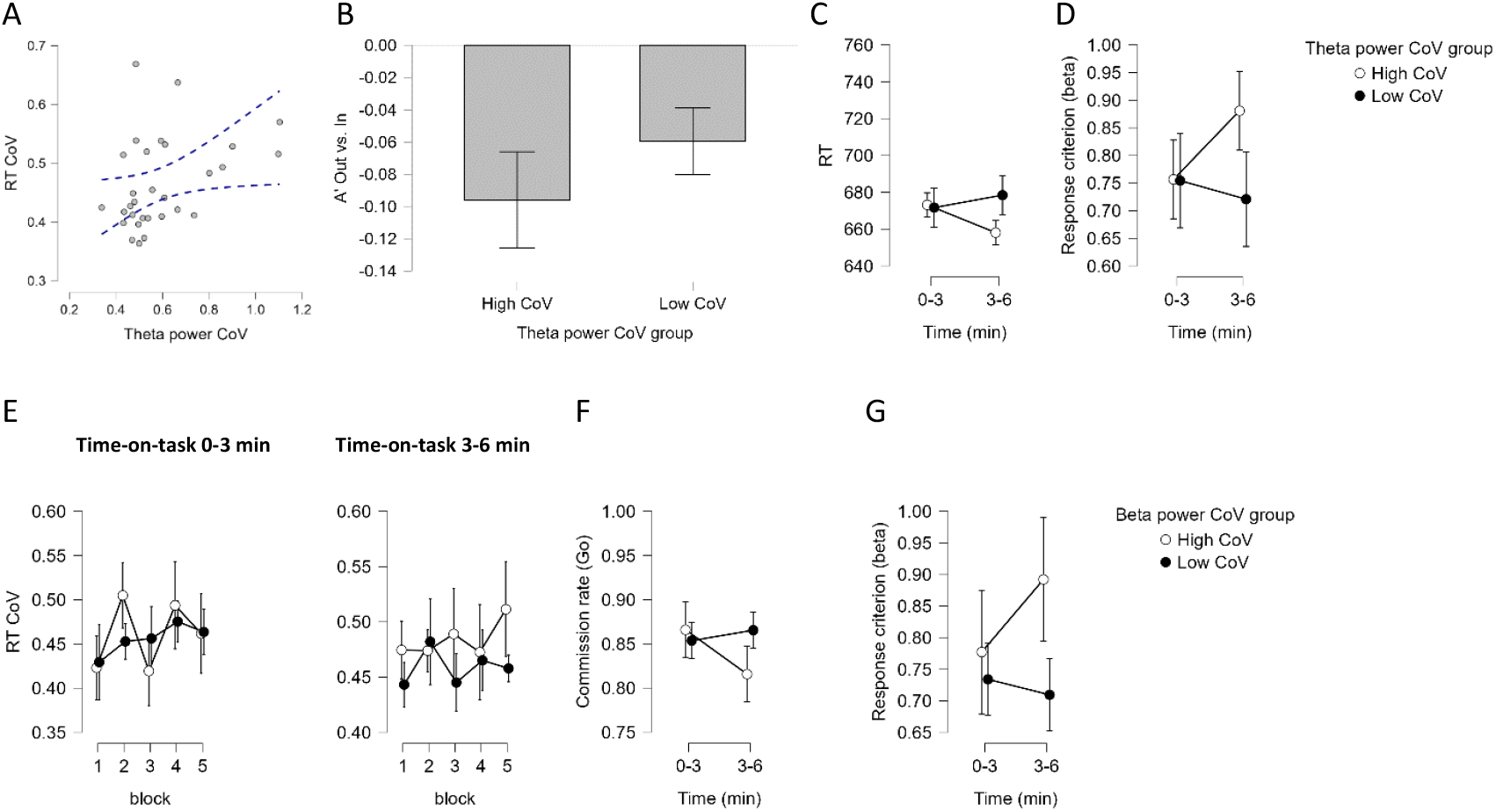
Theta and beta power variability and task performance. (A) Theta power CoV was positively associated with RT CoV. (B) Participants with high theta power CoV showed a stronger decline in sensitivity (A’) during out-versus in-the-zone states, (C) unstable RT with time-on-task, and (D) unstable response bias with time-on-task. (E) Participants with high beta power CoV showed unstable RT CoV across blocks during the first halves of time-on-task (0-3 min, p <.05), (F) a decline in the rate of correct commissions (Go trials) with time-on-task, and (G) unstable response bias with time-on-task.

However, the median split comparisons of groups with high versus low theta power CoV revealed higher differences in A’ between out- and in-the-zone states in the group of high theta CoV compared to those with low theta CoV (t(28)=2.15, p=.04, Cohen’s d=.79; Figure 5B).

In addition, an interaction of group*time-on-task in mean RT (F(1,27)=6.98, p=.014, n^2^_p_=.21), was found in the RT mixed model ANOVA, which reflected that the group of participants with high theta CoV had reduced RT during the second half of the blocks compared to the first half (MDiff=-16 ms, p=.016, Cohen’s d=-.29), while participants with low theta CoV showed no significant differences (MDiff=-6.79 ms, p>.2, Cohen’s d =-.13; Figure 5C). For the mixed model ANOVA on A’, no significant main effects or interactions were found (all p >.17). For the ANOVA on response bias, a significant interaction of group*time-on-task (F(1,27)=4.63, p=.04, n^2^_p_ =.14) revealed that participants with high theta CoV had higher bias during the first halves of time-on-task compared to the second halves (MDiff=-.12, p=.02, Cohen’s d=-.28), while there were no significant differences in the group with low theta CoV (MDiff=.03, p=.52, Cohen’s d=.08; Figure 5D). The block factor did not show a significant interaction with group (high or low theta power CoV) in any of these analyses (all p>.45).

For CoV of beta power around 400 ms after the transition point, a significant interaction of group*block*time-on-task was found in the ANOVA on RT CoV (F(4,108)=4.64, p=.004, n^2^_p_ =.15). The posthoc analysis indicated that participants with high beta CoV showed unstable RT CoV across the different blocks during the first halves of time-on-task (0-3 min, p <.05), but not in the second halves (3-6 min), while no significant differences were found in participants with low beta CoV for time-on-task, or block (all p>.9; Figure 5E). In addition, the ANOVA on commission rates revealed a significant interaction of group*time-on-task (F(1,27)=8.54, p=.007, n^2^_p_ =.24) which reflected that participants with high beta CoV showed a significant decrease of correct commissions rate during the second half of the blocks (MDiff=-.06, p=.002, Cohen’s d=.43), while participants with low beta CoV showed no significant differences (MDiff=.01, p>.2, Cohen’s d=.08; Figure 5F). For the ANOVA of response bias, a trend for a significant interaction of group*time-on-task (F(1,27)=3.49, p=.07, n^2^_p_ =.11) revealed that participants with high beta CoV had higher bias during the first halves of time-on-task compared to the second halves (MDiff=-.11, p=.04, Cohen’s d=-.26), while there were no significant differences in the group with low beta CoV (MDiff=.02, p=.64, Cohen’s d=.06; Figure 5G). No significant main or interactive effects were found in the ANOVA of mean RT. Again, the block factor did not show a significant interaction with the group (beta power CoV) in any of these analyses (all p>.57).

Similar ANOVA comparisons of behavioural measures between groups of high versus low power differences between zones in theta and beta did not yield any significant results (all p>.2).

### Phase alignment across trials

We did not find significant differences between states of in and out-of-the-zone for phase alignment (ITPC) for theta, alpha, or beta oscillations (all cluster statistics =<8, all p>=.23). Therefore, no further analyses were run.

### EEG predictors of performance

Table 1 shows the results for each of the three multiple regression models exploring the associations of theta and beta power decreases in out-of-the-zone (vs. in-the-zone) states, as well as theta power CoV, with behavioural measures. The regression model on RT CoV indicated that theta power decline and theta CoV were the best predictors (β=-.53, t=-3.13, p=.004 and β=.33, t=1.90, p=.069, respectively). Theta power decline in out- (vs. in-) the-zone states was also the best predictor of correct omissions rate (β=.43, t=2.31, p=.029). Finally, Table 1 also shows that beta power changes between zones did not significantly predict any behavioural measure.

**Table 1.**
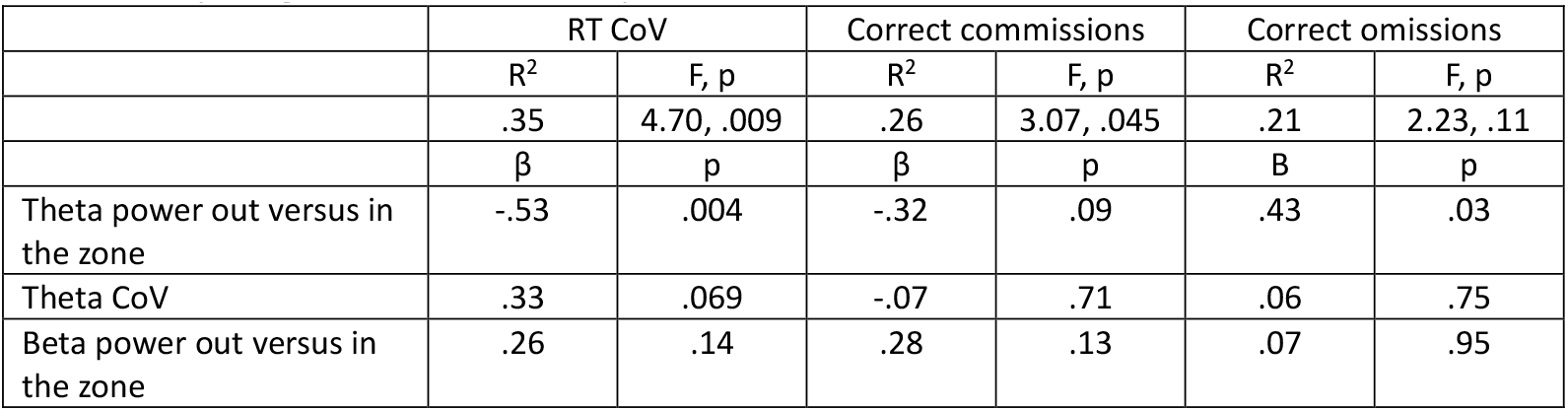
Multiple regression models of EEG predictors of behavioural measures of sustained attention states.

## DISCUSSION

We identified the electrophysiological changes during fluctuating states of sustained attention, using the gradCPT task, which has been shown to be sensitive to reveal sustained attention states of ‘in-the-zone’ or ‘out-of-the-zone’ throughout task performance. Our findings reveal changes in theta and beta power, as well as their power variability, as sensitive neural markers of states of sustained attention and establish also their relation to intra- and inter-individual differences in task performance, while highlighting the critical time windows for the effects in relation to task performance. We discuss each of these findings next.

### Theta power

Our results indicated a transient power reduction of midfrontal theta activity in peri-stimulus transition periods, during out-of-the-zone (as compared to in-the-zone) sustained attention state. The source analysis of this reduction indicated prefrontal and cingular regions, with the maximum difference at medial frontal cortex, corresponding to cognitive control areas (Widge et al. 2019).

The magnitude of the theta power decline during out-of-the-zone (compared to in-the-zone) states was positively associated with increased RT variability, as well as increased commission rates on NoGo trials (i.e. attention lapses). Larger theta power decline during out-of (vs. in-) the-zone states was also found to be specifically associated with a greater difference in attention lapses between out-of-the-zone states compared to in-the-zone states. A larger power decline during out-of-the-zone (vs. in-the-zone) states also correlated with a lower response criterion (larger bias to make a response irrespective of whether the trial is Go or NoGo). This is further suggestive that theta power decline in out-of-the-zone states is indicative of the general reduction in the level of attentional control over performance. Altogether, these findings provide strong support for the role of midfrontal theta oscillations in cognitive control over states of sustained attention, demonstrating a direct association between theta power decline and states of being out-of-the-zone, with the magnitude of this momentary decline specifically predictive of the effect of reduced attention state on task performance. In addition, the variability of theta power throughout the task was also found to predict response variability: higher variability in theta power was associated with higher response variability. Finally, participants with high theta variability also showed a larger difference in their perceptual sensitivity between out-versus in-the-zone states. Overall, theta power in the midfrontal regions (as shown with the source analysis) was revealed as a sensitive marker of sustained attention states and their associated effects on performance.

These results are consistent with a body of findings demonstrating the role of midfrontal theta power in cognitive control over attention. Theta activity is increased in midfrontal regions during effective performance in continuous tasks (Wascher et al. 2014) and during those that require inhibitory control over a conflicting response (Oehrn et al. 2014). Midfrontal theta power also increases after negative feedback during reinforcement learning (van de Vijver et al. 2011) and after conflict detection (Oehrn et al. 2014), as well as after errors of commission (Cavanagh et al. 2012), promoting attentional control reset. Overall, there is a general consensus that midfrontal theta serves as a neurophysiological marker of cognitive control (Cavanagh and Frank 2014; Umemoto et al. 2019), and that specific patterns of theta connectivity and amplitude can reflect specific features of cognitive control (Cooper et al. 2015; Messel et al. 2021). Our findings provide a new demonstration that midfrontal theta power is specifically associated with sustained attention states throughout task performance, and that both theta power and its variability are predictive of performance overall.

We note that some sustained attention studies have previously reported an increase in theta power associated with worse performance as a function of time-on-task (Lal and Craig 2002). In addition, increased theta has been identified as a predictor of attentional decline, including lapses in vigilance and a higher error rate (Boksem et al. 2005; W. Klimesch 1999; Paus et al. 1997; Peiris et al. 2006; Wascher et al. 2014). Indeed, some prior studies have suggested that increased theta power can be considered as a direct index of mental fatigue (Boksem et al. 2005).

While at first sight these findings appear generally inconsistent with the suggestion that increased theta power reflects enhanced attention, this inconsistency can be reconciled by realizing that the studies that linked theta with mental fatigue reported its well-known increase of power with time-on-task as a general, unspecific marker along the scalp. This general increase was reported after apriori averaging across all the EEG channels (e.g. (Lal and Craig 2002)) or even after statistically controlling for scalp regions (Boksem et al. 2005; Wascher et al. 2014). This general increase might reflect a general slowing of oscillatory dynamics due to time-on-task, since additional increases of lower alpha and delta oscillations can be observed as well. On the other hand, the proposal that theta oscillations from midfrontal brain regions does indicate cognitive control as reviewed in Clayton et al. (2015), and including those showing an increase in midfrontal power as a result of time-on-task, relies on the notion that cognitive control is important, not just for achieving effective performance, but also for monitoring attention and its failures and to counteract mental fatigue due to time-on-task (Umemoto et al. 2019). Thus, the increase in midfrontal theta power and the associated increase in rates of errors with a longer time-on-task may reflect the higher demand on task monitoring (especially following errors) during states or reduced attention. This interpretation is supported by findings showing increases of power both in frontal total theta with time-on-task and frontal induced theta during successful resolution of stimulus-response mapping conflict, though we note that the findings fell short of demonstrating an interaction of time-on-task and theta power induced by conflict resolution (Wascher et al. 2014). Similarly, Chidharom et al. (2021) findings of a general increase of midfrontal theta power following errors of commission in a SART type sustained attention task may be taken to reflect the increased monitoring demand following these errors. Moreover, their findings that this effect was less strong in trials with high RT variability (out-of-the-zone states in our study), especially for participants with high RT variability, is in line with our suggestion that midfrontal theta power reflects attentional control over the task, and its decline in intra-individual states of out-of-the-zone, as well as in individuals more prone to out-of-the-zone states, reflect the decline in cognitive control over sustained attention. Our findings are also consistent with those showing that relative increases of midfrontal theta power at the beginning and the end of a 2-hour repetitive time estimation task predict a more accurate and less variable performance (Umemoto et al. 2019).

In addition, our findings of theta power association with sustained attention states throughout the task, rather than only at the later task period, further suggests that periodic increases in theta power are associated with increased cognitive control during in-the-zone states. Conversely, periodic declines in theta power are reflective of reduced cognitive control of attention during out-of-the-zone states, as they occur throughout the task, rather than as a tonic consequence of increased mental fatigue with time-on-task.

Our findings that the change in theta power in relation to attentional states did not occur throughout the entire trial, but at a ∼400 ms period around the transition point between stimuli further suggests that midfrontal theta synchronization can adapt to critical periods where focused attention is particularly needed to perceive the (gradual) transition to a new stimulus. We note that due to the gradual nature of image transitions, the gradCPT paradigm is likely to involve a gradual expectancy build up, once the trial image has been discriminated and responded to, in preparation for the gradual onset of a new image. This process is expected to start following the midpoint of an 800 ms trial, where the image coherence of the current trial image (Nth) starts to be greater than that of the previous trial’s image (Nth-1) and terminates once the Nth image has enough build-up of coherence to be detected. During in-the-zone states, theta synchronization should allow for enhanced attention recruitment during this critical period, which declines with a loss of attention focus (out-of-the-zone states). This aligns with other studies demonstrating the adjustment of the amplitude of oscillations within fine temporal windows during relevant periods for attentional facilitation or blocking of distractors (M. Bonnefond and Jensen 2012; Solis-Vivanco et al. 2018, 2021).

Finally, we note that while our findings point to the role of midfrontal theta in cognitive control over sustained attention, previous work on the rhythmic attention sampling in the theta frequency band during cue-to-target periods in spatial attention tasks has pointed to frontoparietal cortex as the source of activity (Helfrich et al. 2018; Raposo et al. 2023). This regional source difference may point to specialized mechanisms of cognitive control that are involved in sustaining spatial orienting of attention (Fiebelkorn et al. 2018), which our task did not particularly involve.

Overall, our results support the perspective of midfrontal theta as an indicator of cognitive control over sustained attention, attributing lower amplitude to periods of being out-of-the-zone, which involve decline in perceptual sensitivity especially in moments when attention focus is critical for perception. Importantly, they demonstrate that midfrontal theta power is a sensitive marker of sustained attention states throughout task performance.

### Theta phase alignment

We did not find that phase alignment between trials (based on ITPC) was different between being out-versus in-the-zone for theta or other frequencies. Phase alignment can occur in anticipation of a stimulus onset, allowing better encoding when it is relevant or suppression when it is an irrelevant distractor (M. Bonnefond and Jensen 2012; Solis-Vivanco et al. 2018). In sustained attention studies that involve a clear stimuli onset, a decrease in midfrontal theta as well as parieto-occipital phase alignment (measured through ITPC) is found both as a function of increased time-on-task and reduced motivation (Reteig et al. 2019). This suggests that reduced sustained attention (due to either factor) results in reduced ability to focus on the points in time that are critical for stimulus processing. With the gradual change that the gradCPT paradigm involves, the critical time windows are more temporally spread, rather than being based on specific time points of abrupt onset, and this may have reduced the sensitivity of our measure of phase locking.

Nevertheless, our findings that midfrontal theta power was reduced around the (wider-spread, rather than locked to stimulus onset) time window of stimulus transition in out-of-the-zone states, along with an increase of theta power variability, are generally consistent with the interpretation of reduced sustained attention involving also reduced ability to focus on critical time windows for stimulus transition. We note also that in other paradigms that require a more continuous (as opposed to time locked to stimulus onset) attention focus, transient, yet not phase-locked, midfrontal theta is also found to positively correlate with optimal versus sub-optimal states of attention. Specifically, Cohen (2016) used a continuous line tracing task, which allowed the participants to monitor their attention based on accuracy-feedback cues (including gradual changes in the size of the disc pointer that was used to trace the line, which adjusted to a larger size upon imprecise tracing in a previous 5 sec period). Interestingly, midfrontal theta power did not only increase after error commissions on an immediate time scale (e.g. hundreds of ms after) but was also correlated with task performance on intermediate (performance on the previous 5 sec period) and long (scale-free, tens of secs) time scales. These findings led the author to propose that reactive cognitive control as expressed by theta is sensitive to fluctuations of sustained attention at several time scales. Thus, similarly to our findings, theta power modulations have also been found to reflect cognitive control over ongoing transient fluctuations of attention focus throughout the task.

### Theta power variability

The variability of midfrontal theta power was found as a predictor of behavioural variability throughout the task across subjects. At the group level, participants with high theta variability showed greater instability in RT, as well as poorer sensitivity when being out-of-the-zone, compared to participants with low theta variability. This confirms that neurophysiological variability can mirror variability of attentional performance (Vanrullen et al. 2011). Neurophysiological variability reflects fluctuations in the local synchronization of neuronal populations, affecting the stability of their functional role. While fMRI studies have shown that the degree of low frequency (.01–.1 Hz) fluctuations of DMN activity correlates with behavioural variability in this low frequency range (but not in a higher or lower one) during CPT tasks (H. Zhang et al. 2022), to our knowledge, this is the first EEG study reporting a direct association between midfrontal theta power variability and behavioural variability along a continuous performance sustained attention task, as well as an association between power variability and discrimination performance during states of being out-of-the-zone.

### Beta power and its variability

Beta power exhibited an increase in amplitude at the transition point of the scenes during Go trials, and a decrease around 400 ms afterwards. This time course and the source localization to left central regions around the primary motor cortex, suggests that these effects can be attributed to motor planning of the responses (Brown 2006; Park et al. 2013), declining during the transition point and rising once motor prepartion is complete. These effects were reduced during states of out-of (vs. in-) the-zone, suggesting reduced response preparation during periods of weak sustained attention. This is consistent with an interpretation of the increased response bias involving also increased rates of errors of commission in NoGo trials during out-of-the-zone states as reflective of inattentive responding. The finding of the association between a stronger effect of out-of-the-zone on the level of beta suppression during the 400 ms following transition point and increased rates of commission responses in Go trials is in further support of such bias. Finally, the finding that participants with low beta variability at the second latency period (400 ms post transition) showed also a smaller effect of time-on-task on their rate of commission responses in Go trials and response bias hints that beta power variability may also reflect sustained attention to response (which declines with longer time-on-task more so for those with higher beta variability). However, as this pattern of findings reflects a specific interaction with time-on-task effects only in between-group comparisons, it is clear that beta power variability cannot be considered as a general marker of sustained attention.

We note that one study suggested that increased inter-regional synchronization of beta activity (15-30 Hz) reflects increased vigilance, leading to enhanced detection sensitivity of relevant stimuli (i.e. reduced attentional blink) (Gross et al. 2004). Nevertheless, those results were found during a dual-target attentional task. Therefore, generalizing those observations to the sustained attention field has limitations. Indeed, when assessing the predictive ability of theta and beta decline and variability on behaviour, we found that only the decrease in theta amplitude and theta variability predicted RT variability. This confirms frontal theta as the best indicator of sustained attention fluctuations, both within and across subjects.

### Alpha power

We did not find effects of the states of attention zone in the alpha band (8-12 Hz), in contrast to the large body of findings of increased alpha power along with a decline in sustained attention (as expressed in increases in RT variability and error rates) and increased mental fatigue induced with a longer time-on-task (A. Bonnefond et al. 2010; Wascher et al. 2014, 2018). Alpha power increases are known to reflect reduced cortical excitability, which can result from the reduced vigilance (and increased fatigue) as the task goes on (Clayton et al. 2015). The recent findings of periods with increased alpha being associated with reduced entrainment to auditory stimuli demonstrate that periods of reduced cortical excitability can also be found throughout a sustained auditory attention task (Kasten et al. 2024), rather than just in the later task periods. This led us to predict that increased alpha should also be found in states of out-of-the-zone that are revealed throughout our task period.

However, our analyses of sustained attention states did not reveal any time-frequency-channel marker of alpha associated with out-of-the-zone (vs. in-the-zone) attentional states and thus no further analyses (including time on task) could be run on this frequency band. We note however that had time-on-task resulted in increased alpha power in our experiment, then this should have led to increased alpha power in the out-of-the zone (vs in-the-zone) states since these were more prevalent in the later task periods (as seen in our behavioural results).

In addition, it is plausible that the motor requirements of the task in our study might have prevented us from finding differential activity of alpha between attention states. For instance, the constant motor response without clear delay periods among trials may have promoted a desynchronization of alpha activity not only at frontal regions (Babiloni et al. 2002) but across more posterior channels (Y. Wang 2025), reducing our ability to identify power modulations in this band. We note however that our findings of reduced visual detection in states of out-of-the-zone, provide a behavioural indication of reduced visual excitability in these periods (See Carmel et al. (2011)).

### Limitations and future research directions

As the first EEG study to examine the oscillatory correlates of sustained attention states in the widely-used gradCPT, it is important to consider which of the task parameters may be necessary to reveal the effects we reported. For example, our use of 90% Go response versus 10% NoGo response may have increased our task sensitivity to reveal a rapid decline in vigilance as a function of time-on-task within as few as 3 minutes periods. A reversed paradigm with 90% NoGo responses versus 10% Go responses is expected to be less sensitive to reveal these effects (see Jun and Lee (2021)). The high response rate may have also contributed to our task sensitivity to reveal findings in the beta power band (with the source located at the pre-motor frontal cortex), since the demand to respond every 800 ms on 90% of the trials might have increased the need to maintain sustained attention to execute motoric responses at this rate. While sustained attention plays an important role in maintaining the required actions, this may have led to a particularly enhanced role of the beta power in sustained attention in our task. It is plausible that with lower response rates oscillations at the beta frequency will play less of a role in determining the states of attentional zone.

This limitation is unlikely to apply to our findings in the theta band power and its variability, given their convergence with the different sustained attention tasks that have been used in previous research. The findings that various behavioural markers were predicted by theta power bolster the functional role of theta activity for sustained attention. The multiple differences between the pattern found in beta power and its correlations with task performance measures and the pattern found in relation to the theta dynamics (at different time windows to those found in the beta band for example) and its association with behavioural measures, also provide discriminant validity in support of the interpretation that the beta and theta dynamics were independent of each other.

A potential limitation of our task concerned our block-by-block administration of the gradCPT (with short 30 s breaks between each block). This may have weakened our methodological power to reveal additional neurophysiological effects of sustained attention, which can develop throughout an experiment period. For example, it may have rendered our task less sensitive to detect any sustained attention effects reflecting reduced vigilance along with increased alpha activity that may develop towards the later part of the experiment.

Another potential limitation concerns the associations between EEG signals and the task performance measure of RT variability. These should be interpreted with caution, since RT variability was used to define the different states of attention zone in our VTC analysis. It is therefore plausible that the different states of attention zone were more distinct in individuals with a larger RT CoV, and this may have contributed (at least in part) to the sensitivity of their data to reveal differences in EEG signals between the states.

Finally, although we have used source analysis to clarify the cortical brain regions that are likely to have generated the EEG activity we found, EEG does not have the sensitivity to reveal differences between specific brain regions, and while the midfrontal cortex activity in the theta oscillations band that was found to reflect states of sustained attention zone in our study may correspond to mPFC involvement in the DMN, this is merely suggestive. A combined fMRI-EEG study might reveal the specific relation of our findings of theta synchronization and attention states to the large body of previous fMRI studies using the gradCPT, which show the involvement of the DMN and DAN networks in these states.

## Conclusions

In conclusion, we found that oscillatory changes in midfrontal theta and prefrontal beta power, and in theta power variability, are associated with different states of sustained attention throughout task performance. Specifically, we show that midfrontal theta synchronization and prefrontal beta desynchronization during critical time periods where focused attention is particularly needed for perception determines being in an effective (in-the-zone) versus ineffective (out-of-the-zone) sustained attention state characterized by increased response variability and reduced perceptual sensitivity. Moreover, our findings demonstrated that a stronger decline in theta power in out-of-the-zone (vs. in-the-zone) states was a predictor of an individual’s level of response variability and ability to accurately inhibit responses upon the appearance of a NoGo target.

The variability of midfrontal theta power during task performance was also found to predict individual differences in various behavioural measures of attentional decline (i.e. RT CoV and decline in sensitivity during out-of-the-zone states), further supporting the proposal that the variability in theta activity indicates reduced ability to sustain a stable level of cognitive control over sustained attention.

Overall, our findings demonstrate that the synchronization dynamics of midfrontal theta and prefrontal beta roscillations in time windows critical for perception, are strong determinants of states of sustained attention throughout a task.

## IMPACT STATEMENT

Our research establishes the neurophysiological correlates of sustained attention states throughout task performance, as they periodically fluctuate from focused ‘in-the-zone’ to ‘out-of-the-zone’ states, prone to attention lapses. The results highlight midfrontal theta oscillations power-specifically during critical time periods for perception-as an index of attentional states and demonstrate that theta power variability can predict task performance measures of attention lapses.

## ACKNOWLEDGMENTS

None.

